# Validation of saline, PBS and a locally produced VTM at varying storage conditions to detect the SARS-CoV-2 virus by qRT-PCR

**DOI:** 10.1101/2022.08.29.505649

**Authors:** Caroline Ngetsa, Victor Osoti, Dorcas Okanda, Faith Marura, Krupali Shah, Henry Karanja, Daisy Mugo, John Gitonga, Martin Mutunga, Clement Lewa, Benedict Orindi, Philip Bejon, Lynette Isabella Ochola-Oyier

## Abstract

Coronavirus Disease-2019 tests require a Nasopharyngeal (NP) and/or Oropharyngeal (OP) specimen from the upper airway, from which virus RNA is extracted and detected through quantitative reverse transcription-Polymerase Chain Reaction (qRT-PCR). The viability of the virus is maintained after collection by storing the NP/OP swabs in Viral Transport Media (VTM).

We evaluated the performance of four transport media: locally manufactured (“REVITAL”) Viral Transport Media (RVTM), Standard Universal Transport Media (SUTM), PBS and 0.9% (w/v) NaCl (normal saline). We used laboratory cultured virus to evaluate: i) viral recovery and maintaining integrity at different time periods and temperatures; ii) stability in yielding detectable RNA consistently for all time points and conditions; and iii) their overall accuracy.

Four vials of SARS-CoV-2 cultured virus (2 high and 2 low concentration samples) and 1 negative control sample were prepared for each media type (SUTM, RVTM, PBS and normal saline) and stored at the following temperatures, -80°C, 4°C, room temperature (25°C) and 37°C for 7 days. Viral Ribonucleic acid (RNA) extractions and qRT-PCR were done on the following days after inoculation with the cultured virus, days 1, 2, 3, 4 and 7 to assess virus stability and viral recovery.

C_T_ values fell over time at room temperature, but normal saline, PBS, RVTM and SUTM all showed comparable performance in maintaining virus integrity and stability allowing for the detection of SARS-CoV-2 viral RNA.

Overall, this study demonstrated that normal saline, PBS and the locally manufactured VTM can be used for COVID-19 sample collection and testing, thus expanding the range of SARS-CoV-2 viral collection media.

## Introduction

During the initial stages of the Coronavirus Disease-2019 (COVID-19) pandemic, caused by the Severe Acute Respiratory Syndrome Coronavirus 2 (SARS-CoV-2), public health authorities across the globe resorted to mass testing as a strategy to contain the fast-spreading disease(1). A standard COVID-19 test requires a Nasopharyngeal (NP) and/or Oropharyngeal (OP) specimen from the upper airway, from which virus RNA is extracted and detected through quantitative reverse transcription-Polymerase Chain Reaction (qRT-PCR)(2,3). Additionally, the accuracy of a test result is dependent on the integrity of the pre-analytical step, which involves sample collection, storage, transport, and maintenance in cold chain. The viability of the virus is maintained after collection by storing the NP/OP swabs, in Viral Transport Media (VTM), and cold chain shipment to the laboratory testing station. Several types of VTM exist and have been used to transport samples for the diagnosis of other infectious agents such the H1N1 strain of Influenza A, Herpes simplex and adenovirus(4). For COVID-19 diagnosis, the Unites States (US) Centre for Disease Control gave recommendations for the components of an appropriate VTM. These include a Sterile Hanks Balanced Salt Solution (HBSS), heat-inactivated Fetal Bovine Serum (FBS) and a suitable antibiotic or anti-fungal agent(5). However, the magnitude of the pandemic necessitated an increase in the testing capacity of diagnostic laboratories, leading to marked disruptions in the supply chain of reagents, including VTM(6).

Different VTM are therefore currently available commercially, including DNA/RNA Shield™, the IMPROVIRAL™ NAT medium, Minimum Essential Medium (MEM), 1X Phosphate-buffered Saline (PBS) and normal saline (0.9% (w/v) NaCl). Although PBS and normal saline are considered non-conventional transport media and may not be optimal for the storage of NP/OP samples(6), they have been shown to perform well as alternatives to the commercially available, mainstream transport media(7,8). For instance, Penrod et al., 2021 evaluated five transport media (VTM (containing 29.5g/l tryptose phosphate, 5g/l gelatin, 10,000U/l penicillin, 10,000U/l streptomycin, 25μg/l Fungizone), Copan Universal Transport Medium™, Becton Dickinson Liquid Amies Elution Swab (Eswab) Collection/Transport System, Remel Microtest™ M4RT® Multi-Microbe Media, and sterile 0.9% (w/v) sodium chloride) and reported that they all had equivalent sensitivity/specificity in COVID-19 diagnosis. Similar findings were obtained by several other studies which confirmed that PBS and normal saline not only maintained the stability and integrity of viral RNA (for up to 1 month when stored at -80ºC and 7 days at 4ºC) but also provided results that matched those obtained from other media(3,7,9,10). On the contrary, a different study demonstrated conflicting results of viral samples in 0.9% (w/v) saline not performing as well as commercially available transport media over a 72-hour storage period, although comparisons in this study were of samples during stock outs of VTM with those where VTM was available, rather than for split or standardized samples(11). Furthermore, a recent report showed that PBS was less sensitive compared to other media such as DNA/RNA shield™, highlighting the need for further research to ascertain the stability of this alternative transport medium(3).

Kenya, like several countries during the COVID-19 pandemic, suffered from disruptions in the supply of reagents required for diagnosis. It was therefore imperative to seek alternatives that would cushion diagnostic centres from unforeseen shortages. Consequently, the present study evaluated the performance of four transport media, i.e., a locally manufactured (i.e. “REVITAL”) Viral Transport Media (RVTM), Standard Universal Transport Media (SUTM), PBS and 0.9% (w/v) NaCl as plausible options for the transportation and storage of NP/OP samples. We used a laboratory cultured virus to evaluate: i) viral recovery and maintaining integrity at different time periods and temperatures; ii) stability in yielding detectable RNA consistently for all time points and conditions; and iii) their overall accuracy. These were all assessed against of the Standard, Universal Transport Media (UTM).

## Methods

### Virus inoculation

Cultured SARS-CoV-2 was used to prepare virus stocks in minimum essential medium (MEM 2%) (Gibco, Thermo Fisher Scientific) containing 2% fetal bovine serum (FBS) (Gibco, United States). We first generated a10-fold dilution series (viral dilutions 1 to 10, S1 Table) that were used to identify the high concentration (virus dilution 3) cycle threshold (Ct) values of 19 and low concentration (virus dilution 6) Ct of 30 (approximately 1,000,000 to 1,000 nucleic acid copies per ml, respectively).

We inoculated the high and low concentration viral dilutions in standard universal transport medium (SUTM, Copan Diagnostics, Italy), a locally produced viral transport medium (REVITAL Healthcare, Kenya) (RVTM), 1X Phosphate Buffered Saline (PBS) (OXOID, Hampshire, England) and 0.9% (w/v) sodium chloride (NaCl) (SIGMA). Five vials (2 high concentration SARS-CoV-2 positive samples, 2 low concentration SARS-CoV-2 positive samples and 1 negative control) were prepared for each media type and stored at the following temperatures, -80°C, 4°C, room temperature (25°C) and 37°C for 7 days (Fig 1), generating a total of 80 vials. Viral Ribonucleic acid (RNA) extractions and qRT-PCR were done on the following days after inoculation with the cultured virus, days 1, 2, 3, 4 and 7 to assess virus stability and viral recovery. All in-house laboratory prepared buffers and media were filtered and autoclaved before use.

**Fig 1.**
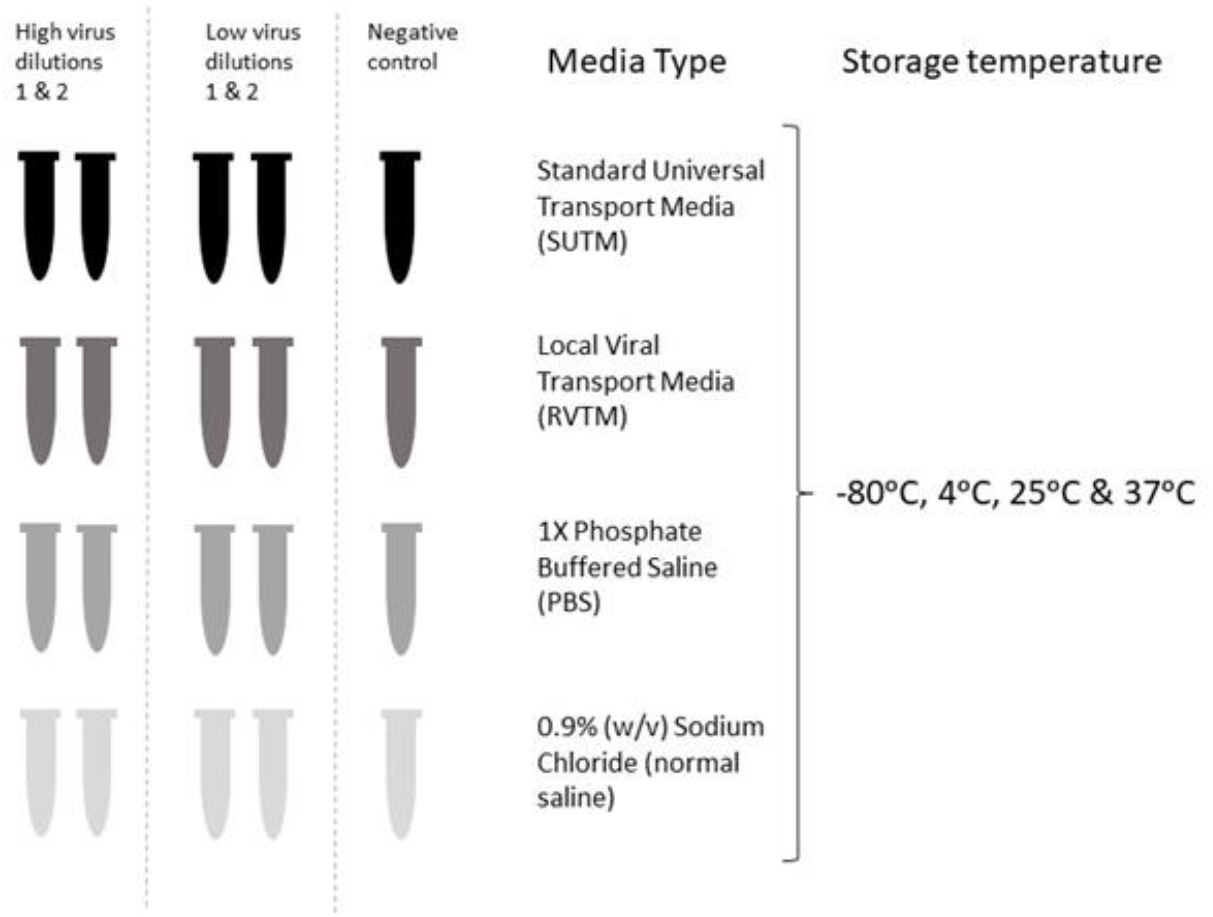
An illustration showing virus dilutions in different media types and different virus inoculation temperatures. Each media type was made up of 2 tubes inoculated with high virus concentrations, 2 tubes of low virus concentrations and 1 negative control tube, generating a total of 5 tubes stored at the following temperatures: -80°C, 4°C, 25°C (room temperature) and 37°C. Thus, each media type consisted of a total of 80 tubes that were subsequently used to extract RNA for qRT-PCR.

### RNA extraction

Viral RNA was extracted from serially diluted and the 4 different media SARS-CoV-2 samples using the QIAamp Viral RNA Mini Kit (QIAGEN GmbH, Hilden, Germany). Briefly, 140μl sample was mixed with 280μl of viral lysis buffer and 2.8μl of carrier RNA. Thereafter, 280μl of absolute ethanol was added and the contents were briefly vortexed before transferring into QIAamp Mini Spin Columns. The spin columns were spun at 6,000 x g for 1 minute, the elute was discarded and 500μl of buffer AW1 was added followed by centrifugation at 6,000 x g for 1 minute.

The flowthrough was discarded and 500μl of buffer AW2 was added followed by centrifugation at 16,000 x g for 3 minutes. The elute was discarded and the spin columns allowed to air dry for 5 minutes before elution with 60μl buffer AVE.

### Quantitative reverse transcription PCR assays

For the qRT-PCR assay, the European Virus Archive – GLOBAL (EVAg) primer/probe set containing 1.75μl primer probe mix, 3.75μl Nuclease free water, 2.5μl TaqMan™ Fast Virus master mix and 2.5μl of template RNA were used(12). Cycling conditions were 50°C for 5min, 95°C for 20sec then 40 cycles of 95°C for 3sec and 58°C for 45sec on the QuantStudio 5 and 7 qRT-PCR systems (Applied Biosystems, CA). A known negative sample and non-template control (NTC) sample were included in each run. All samples were done in duplicates.

Ct values obtained for each sample were analyzed to assess the impact of the four media types and storage condition on Ct values. A test was considered successful if the negative and NTC samples did not have Ct values. Furthermore, data obtained from PCR analyser were in Ct values. These were reviewed and interpreted using a Ct cut-off of 36, interpreting those values of less than and greater than the cut-off as positive and negative test results, respectively. The qRT-PCR machine also gave an ‘undetermined’ result among the Ct values, these were interpreted as negative test results.

### Statistical analysis

All analyses were performed using Stata version 15 (Stata Corp., College Station, TX). Graphical presentations were produced in R version 3.6.1(13). First, we note that there were 128 negative samples defined as “undetermined” that were replaced with an arbitrary high Ct for negative (i.e., 37) to uncover the bias that might occur due to their exclusion during statistical analysis. Temporal (in days) and temperature trends on viral detection across the four media types were assessed. A Spearman’s correlation coefficient was obtained between the SUTM and the other 3 media types and a line of best fit was estimated using a Deming regression model(14). Accuracy of the media types was determined by estimating the positive percent agreement (PPA), negative percent agreement (NPA), and overall percent agreement (OPA)(15). Finally, Ct values were compared using a multiple linear regression model with main effects for media type, temperature, concentration (low/high/negative control), day and the interactions for media type and day, and temperature and day as covariates. Overall effects were tested using the Wald test. All tests were performed at a 5% significance level.

## Results

### qRT-PCR

In total, 811 qRT-PCR assays were conducted for all 4 media types at all 4 temperatures over a 7-day period. A total of 15 negative controls were incorporated for all the run days and these yielded negative results as expected. However, there were 16 (1.98%) negative samples with Ct<36. These were regarded as being contaminated and hence excluded from any further analysis. They were primarily observed at -80^0^C and 4^0^C across all media types (S2 Table and Fig 2). Of the remaining 780 samples there were 194 (24.9%) for SUTM, 195 (25%) for PBS, 193 (24.7%) for RVTM, and 198 (25.4%) samples for normal saline. Of these, 640 (82%) gave positive test results while 140 (18%) gave negative test results as expected.

**Fig 2.**
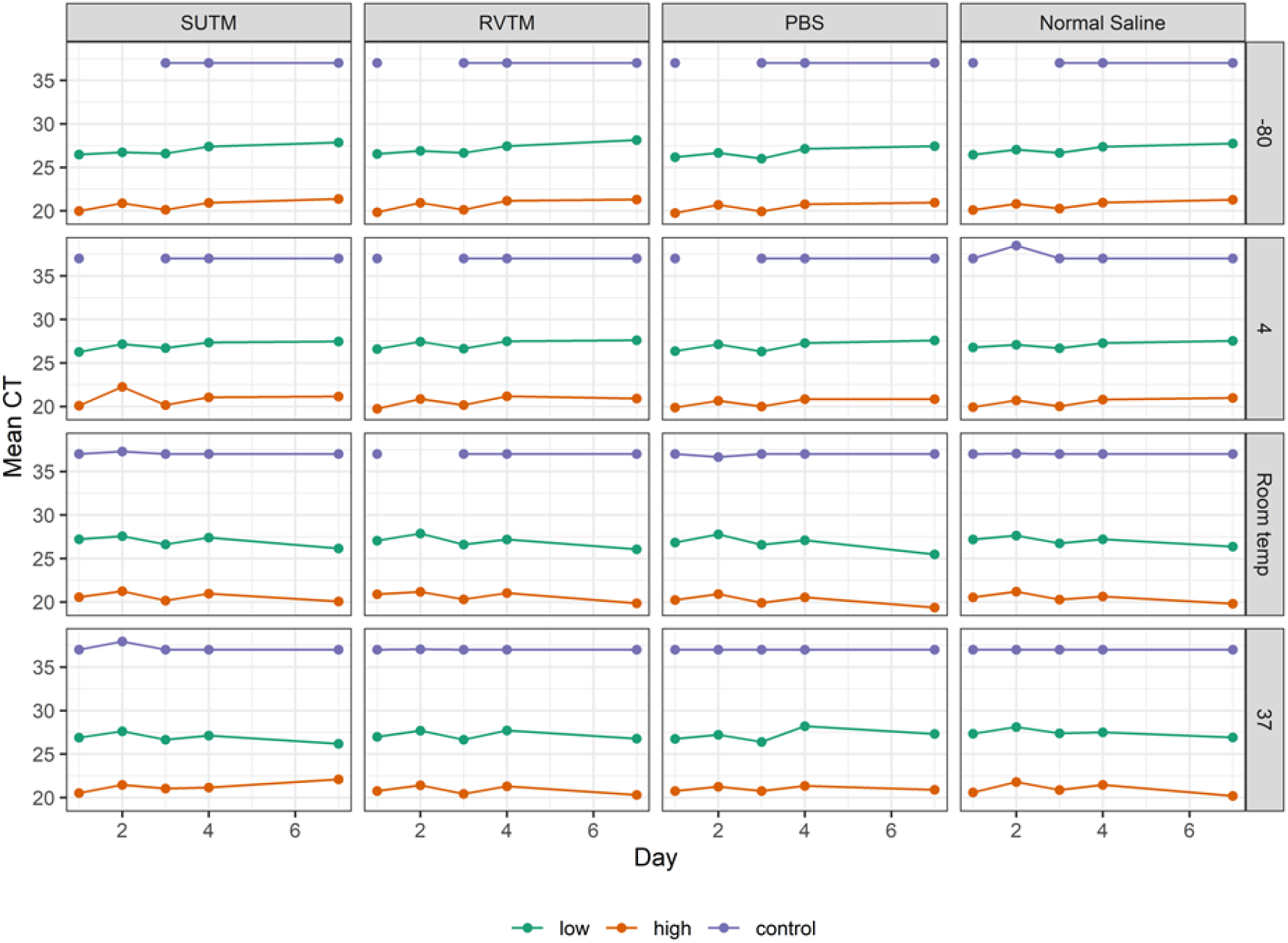
Mean Ct values for the four media types over time (in days) at -80°C, 4°C, room temperature (25°C) and 37°C stratified by concentration. A Ct>36 or 37 indicates a negative result. The gaps in temperatures -80°C and 4°C indicate contaminated negative samples, which were excluded from the analyses.

### Comparisons between the 4 media types

For all temperature levels, media types and days the Ct values were consistently highest in the low concentration samples (between Cts 25 and 28) and they were significantly higher than those for high concentration samples (between Ct 18 and 23) (Tables 1 and 2, Fig 2). The Ct values varied over time in the four temperature levels examined (Table 2). Ct values generally increased over time (day) in the media stored at -80ºC and 4ºC, decreased and remained the same at room temperature and 37ºC, respectively (Fig 2). The multiple linear regression analysis demonstrated there were no statistically significant differences in Ct values across the media types (Table 2; Fig 1). The correlations of Ct values between the 4 media types were generally higher at low than high concentrations (Figs 3A and 3B). The overall accuracy of RVTM, PBS and saline compared to SUTM was 100% (Table 3).

**Table 1.** Mean Ct values by media, temperature, and time (in days) separately for high and low concentrations, and negative controls

|  | Concentration |  | Negative controls<br>(N=140) |
| --- | --- | --- | --- |
|  | Low<br>(N=320) | High<br>(N=320) |  |
| Media |  |  |  |
| SUTM | 26.97 (0.53) | 20.86 (0.91) | 37.07 (0.35) |
| RVTM | 27.10 (0.57) | 20.68 (0.57) | 37.02 (0.10) |
| PBS | 26.89 (0.68) | 20.51 (0.64) | 36.98 (0.12) |
| Normal Saline | 27.15 (0.51) | 20.66 (0.54) | 37.08 (0.35) |
| Temperature |  |  |  |
| -80 | 26.97 (0.60) | 20.61 (0.55) | 37.00 (0.00) |
| 4 | 27.04 (0.47) | 20.60 (0.69) | 37.08 (0.37) |
| Room temperature | 26.93 (0.63) | 20.49 (0.54) | 37.02 (0.21) |
| 37 | 27.17 (0.60) | 21.02 (0.82) | 37.05 (0.29) |
| Concentration |  |  |  |
| Low | 27.03 (0.58) |  |  |
| High |  | 20.68 (0.69) |  |
| Control |  |  | 37.04 (0.26) |
| Days |  |  |  |
| 1 | 26.74 (0.41) | 20.26 (0.47) | 37.00 (0.00) |
| 2 | 27.35 (0.46) | 21.14 (0.57) | 37.30 (0.69) |
| 3 | 26.62 (0.33) | 20.28 (0.41) | 37.00 (0.00) |
| 4 | 27.38 (0.32) | 21.00 (0.36) | 37.00 (0.00) |
| 7 | 27.04 (0.80) | 20.71 (0.94) | 37.00 (0.00) |

**Table 2.**
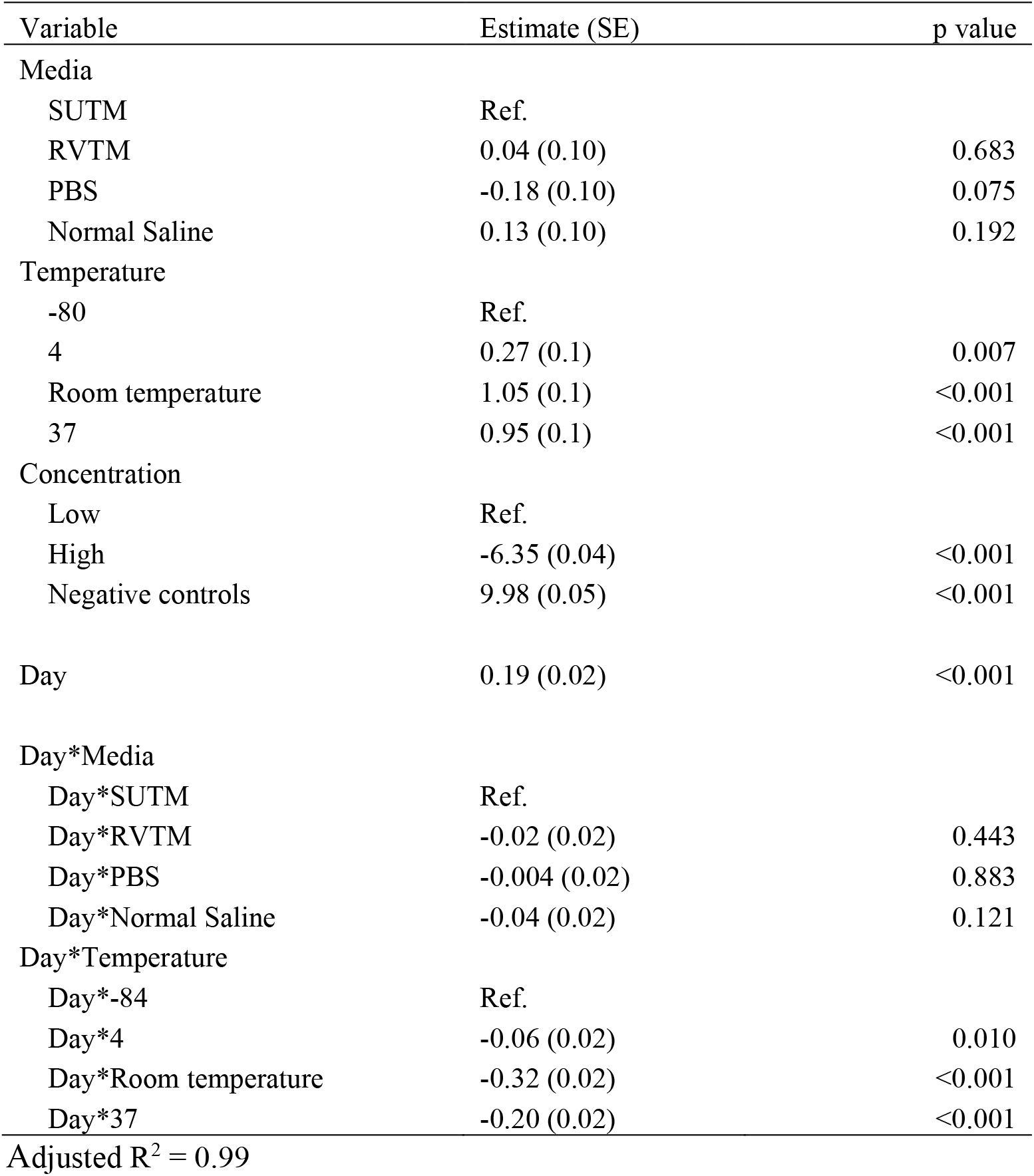
Multiple linear regression analysis to compare the differences between the media

**Table 3.** Overall agreement between RVTM and SUTM for all time points and conditions*

|  | **SUTM (Reference)** |  |  |  |  |
| --- | --- | --- | --- | --- | --- |
| **RVTM (Comparator)** | Positive | Negative | PPA | NPA | OPA |
| Positive | 160 | 0 |  |  |  |
| Negative | 0 | 33 |  |  |  |
| Total | 160 | 33 | 100 (97.6-100) | 100 (89.6-100) | 100 (98-100) |
| **PBS (Comparator)** |  |  |  |  |  |
| Positive | 160 | 0 |  |  |  |
| Negative | 0 | 34 |  |  |  |
| Total | 160 | 34 | 100 (97.7-100) | 100 (89.9-100) | 100 (98.1-100) |
| **Saline (Comparator)** |  |  |  |  |  |
| Positive | 160 | 0 |  |  |  |
| Negative | 0 | 35 |  |  |  |
| Total | 160 | 35 | 100 (97.7-100) | 100 (90.1-100) | 100 (98.1-100) |

PPA=Positive percent agreement; NPA=Negative percent agreement; OPA= Overall percent agreement

**Fig 3.**
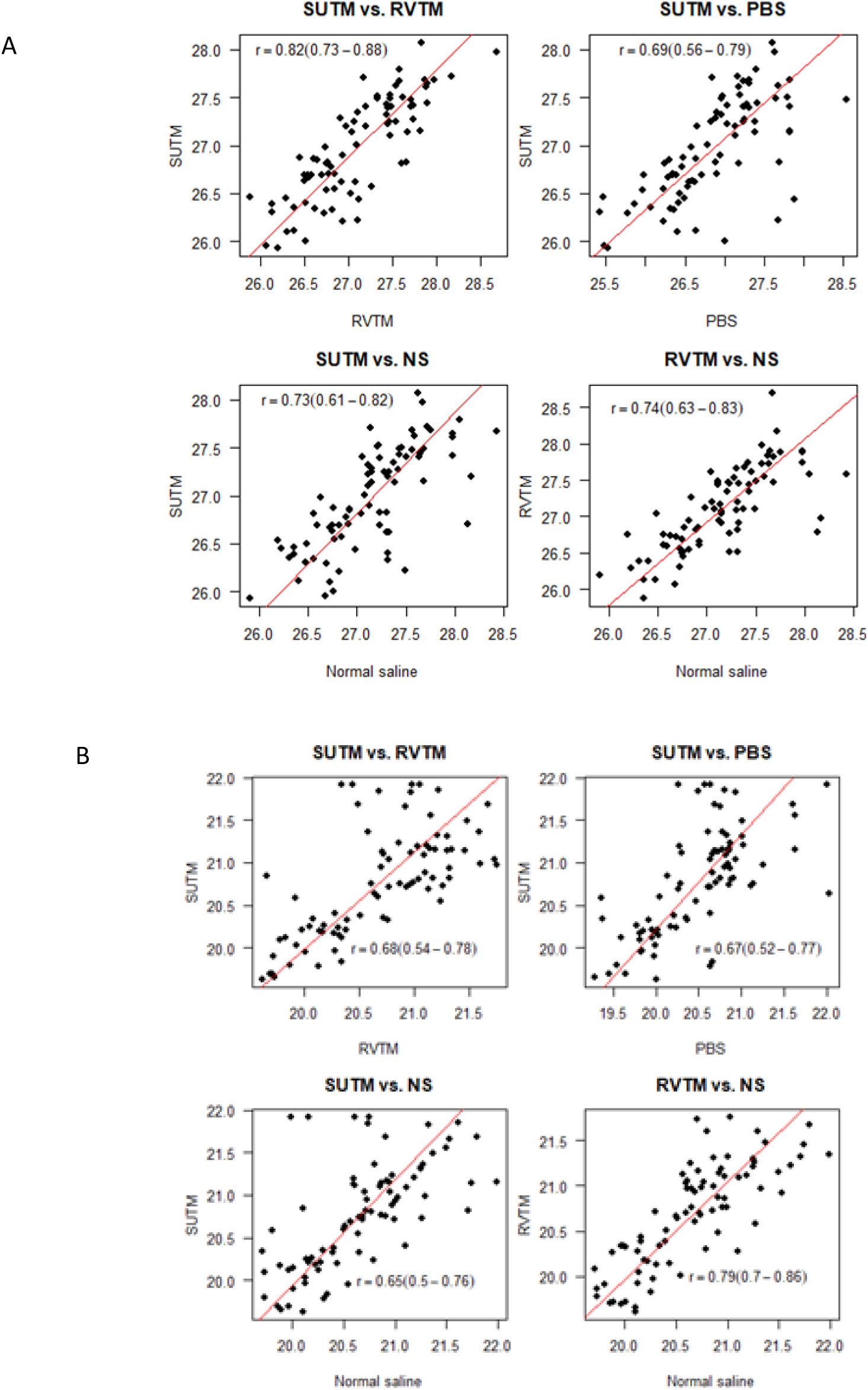
Correlations between Ct values from the 4 media types. **(A)** Scatter plots for Ct values at low concentration. **(B)** Scatter plots for Ct values at high concentration. The line of best fit was estimated using a Deming regression model.

## Discussion

Normal saline, PBS and the locally manufactured VTM showed comparable performance in maintaining virus integrity and stability allowing for the detection of SARS-CoV-2 viral RNA. All the media types supported the recovery of SARS-CoV-2 to as low as 1000 viral nucleic acid copies per ml. Furthermore, even at temperatures above 25°C virus was detected over the 7-day sampling frame.

There were variations in RNA yield based on temperature storage conditions and day of testing, notably in the samples stored at -80°C and 4°C, which showed a reduction in viral load over time. Similar findings have been reported previously, including in other respiratory viruses such as Herpes simplex, Influenza, enterovirus and adenovirus(4,10). Viral recovery appeared more stable at 37°C than at room temperature. Additionally, normal saline appeared to have lower Ct values post-day 4 at 37°C compared to SUTM. A previous study described better sensitivity to detect Ct values in samples in saline across 4, 25 and 37°C, low Ct values were also detected in samples stored at 37°C(16). These observations are not unusual since RNA is an unstable molecule that degrades over time and in different temperature conditions(17). The reliability of Ct results on an assessment of the media types revealed consistency at high RNA concentrations. This is expected, since at low viral loads there is an increased signal to noise ratio as the limit of detection of the qRT-PCR test is reached and taking into consideration the variability between qRT-PCR kits.

There were some contaminated negative control samples (Ct<36) in <2% of the samples examined. The contamination may have occurred during extraction or the compilation of qRT-PCR reagents(18) due to amplicon carry-over given the high number of SARS-CoV-2 amplicons being generated in the lab. It may also be a reflection of the high signal-to-noise ratio at high Ct values. We had mitigated against contamination by incorporating proper disinfection, and use of dedicated working spaces ensuring minimal viral contamination(18). Samples paired with positive negative controls were regarded as unreliable and were removed from further analyses.

These results based on laboratory cultured samples stored at a range of temperatures suggest that samples swabbed directly from patients could be transported in the different media types examined in this study, given the comparable ability to detect the viral RNA. However, we did not test whether the different media types support viable culturable virus.

The cultured virus we used may be at higher concentration than samples taken in the field, and storage conditions during transport may be more variable. However, we did not observe evidence of any differences in RNA levels based on the PCR results by VTM, including when virus was stored at room temperature.

## Acknowledgements

We thank Prof George Warimwe’s group from KEMRI-Wellcome Trust Research Programme for the cultured SARS-CoV-2 virus aliquot they kindly provided for use in this study. We thank the KEMRI Director for permission to publish this work.

## Supporting Information

**S1 Table**. SARS-CoV-2 cultured virus dilution series and corresponding C_T_ values

**S2 Table**. Numbers of negative samples with a C_T_<36 for each media type and at each temperature tested

## References

1. Shen M, Xiao Y, Zhuang G, Li Y, Zhang L. Mass testing—An underexplored strategy for COVID-19 control. The Innovation. 2021 May 28;2(2).

2. Kevadiya BD, Machhi J, Herskovitz J, Oleynikov MD, Blomberg WR, Bajwa N, et al. Diagnostics for SARS-CoV-2 infections. Vol. 20, Nature Materials. Nature Research; 2021. p. 593–605.

3. Barrera-Avalos C, Luraschi R, Vallejos-Vidal E, Figueroa M, Arenillas E, Barría D, et al. Analysis by real-time PCR of five transport and conservation mediums of nasopharyngeal swab samples to COVID-19 diagnosis in Santiago of Chile. J Med Virol. 2022 Mar 1;94(3):1167–74.

4. Druce J, Garcia K, Tran T, Papadakis G, Birch C. Evaluation of swabs, transport media, and specimen transport conditions for optimal detection of viruses by PCR. J Clin Microbiol. 2012 Mar;50(3):1064–5.

5. CDC. Centers for Disease Control and Prevention PREPARATION OF VIRAL TRANSPORT MEDIUM [Internet]. 2020 [cited 2022 May 10]. Available from: https://www.google.com/search?q=components+of+VTM%2C+CDC&rlz=1C1GCEA_enKE999KE1000&oq=components+of+VTM%2C++CDC&aqs=chrome..69i57.7630040j0j1&sourceid=chrome&ie=UTF-8

6. Penrod Y, Garcia D, Dunn ST. Evaluation of transport media for laboratory detection of SARS-CoV-2 in upper respiratory tract swab specimens. J Med Virol. 2021 May 1;93(5):2774–81.

7. Rodino KG, Espy MJ, Buckwalter SP, Walchak RC, Germer JJ, Fernholz E, et al. Evaluation of saline, phosphate-buffered saline, and minimum essential medium as potential alternatives to viral transport media for SARS-CoV-2 testing. Vol. 58, Journal of Clinical Microbiology. American Society for Microbiology; 2020.

8. Borkakoty B, Jakharia A, Bali N, Sarmah M, Hazarika R, Baruah G, et al. A preliminary evaluation of normal saline as an alternative to viral transport medium for COVID-19 diagnosis. Vol. 153, Indian Journal of Medical Research. Wolters Kluwer Medknow Publications; 2021. p. 684–8.

9. Amy A Rogers, Russell E Baumann, Gwynngelle A Borillo, Ron M Kagan, Hollis J Batterman, Marzena M Galdzicka, et al. Evaluation of Transport Media and Specimen Transport Conditions for the Detection of SARS-CoV-2 by Use of Real-Time Reverse Transcription-PCR. J Clin Microbiol. 2020 Jul 23;58(8):1–5.

10. Garrett A. Perchetti, Meei-Li Huang, Vikas Peddu, Keith R Jerome, Alexander L Greninger. Stability of SARS-CoV-2 in Phosphate-Buffered Saline for Molecular Detection. J Clin Microbiol. 2020 Aug;58(8):1–2.

11. Garnett L, Bello A, Tran KN, Audet J, Leung A, Schiffman Z, et al. Comparison analysis of different swabs and transport mediums suitable for SARS-CoV-2 testing following shortages. J Virol Methods. 2020 Nov 1;285.

12. Mohammed KS, de Laurent ZR, Omuoyo DO, Lewa C, Gicheru E, Cheruiyot R, et al. An optimisation of four SARS-CoV-2 qRT-PCR assays in a Kenyan laboratory to support the national COVID-19 rapid response teams. Wellcome Open Res. 2020 Jul 7;5:162.

13. R Core Team. R: A language and environment for statistical computing. R Foundation for Statistical Computing, Vienna, Austria. [Internet]. 2019 [cited 2022 Jul 21]. Available from: https://www.R-project.org/

14. Henk Konings. Use of deming regression in method-Comparison studies. Survey of Immunologic Research. 1982 Dec;1:371–4.

15. Clinical and Laboratory Standards Institute. User Protocol for Evaluation of Qualitative Test Performance, 2nd Edition [Internet]. 2008 [cited 2022 Jul 21]. Available from: https://clsi.org/standards/products/method-evaluation/documents/ep12/

16. Sinha P, Jain DK, Gupta S, Gupta M, Gupta M, Agarwal A, et al. Evaluation of Saline as Potential Alternative to Viral Transport Media for COVID-19 Samples Stored at Different Temperatures. JOURNAL OF CLINICAL AND DIAGNOSTIC RESEARCH. 2021;

17. Fabre AL, Colotte M, Luis A, Tuffet S, Bonnet J. An efficient method for long-term room temperature storage of RNA. European Journal of Human Genetics. 2014 Mar;22(3):379–85.

18. Borst A, Box ATA, Fluit AC. False-positive results and contamination in nucleic acid amplification assays: Suggestions for a prevent and destroy strategy. Vol. 23, European Journal of Clinical Microbiology and Infectious Diseases. 2004. p. 289–99.

