## Supplementary Tables for "Validation of saline, PBS and a locally produced VTM at varying storage conditions to detect the SARS-CoV-2 virus by qRT-PCR"

**S1 Table.**

| Virus dilution | Dilution | RADI_RdRp | RADI_S_GENE |
| --- | --- | --- | --- |
| 1 | neat | 11.85 | ND |
| 2 | 10^-1^ | 13.17 | 14.73 |
| 3 | 10^-2^ | 15.65 | 18.21 |
| 4 | 10^-3^ | 18.9 | 21.4 |
| 5 | 10^-4^ | 23.3 | 25.82 |
| 6 | 10^-5^ | 26.27 | 28.54 |
| 7 | 10^-6^ | 29.78 | 32.48 |
| 8 | 10^-7^ | 33.64 | 35.89 |
| 9 | 10^-8^ | 37.27 | 37.15 |
| 10 | 10^-9^ | Undetermined | Undetermined |
| 11 | 10^-10^ | Undetermined | Undetermined |

**S2 Table.**

|  | Temperature | | | |  | |
| --- | --- | --- | --- | --- | --- | --- |
| Media | -80°C | 4°C | Room  temperature | 37°C | | Total |
| SUTM | 4 | 1 | 0 | 0 | | 5 |
| RVTM | 2 | 2 | 1 | 0 | | 5 |
| PBS | 2 | 2 | 0 | 0 | | 4 |
| Normal Saline | 2 | 0 | 0 | 0 | | 2 |
| Total | 10 | 5 | 1 | 0 | | 16 |
